# CRISPR-based neuromorphic computing for solving regression and classification

**DOI:** 10.64898/2025.12.03.692209

**Authors:** Claudia Montufar Leon, Yao Wang, Frank Britto Bisso, Arya Mehta, Christian Cuba Samaniego

## Abstract

The CRISPR-dCas9 system has emerged as a versatile platform for programmable gene regulation, offering unique advantages in modularity and orthogonality for constructing synthetic genetic circuits. Here, we present a novel architecture for biomolecular neural networks based on dCas9, guide RNAs, and antisense RNA sequestration. Through mathematical modeling and steady-state analysis, we demonstrate that this system functions as a molecular perceptron with a threshold activation function analogous to a saturated rectified linear unit (ReLU). However, a critical challenge in scaling these circuits is competition for the finite dCas9 pool, whose expression must remain low to avoid cytotoxicity. We address this constraint by developing a resource-aware design framework and characterizing how shared dCas9 availability affects network performance. Our results show that for classification tasks, decision boundaries remain invariant under resource competition, while for regression tasks, node thresholds are preserved despite sensitivity in output magnitude under heterogeneous binding conditions. We demonstrate the computational capabilities of this platform through both linear and nonlinear classification problems, as well the approximation of a band-pass function as a proof-of-concept regression task. This work expands the repertoire of molecular mechanisms capable of computation and establishes design principles for implementing CRISPR-based neuromorphic circuits that can execute complex computational tasks within the biochemical constraints of living cells.

## I. Introduction

The CRISPR–Cas9 system has rapidly evolved from a bacterial adaptive immune mechanism into one of the most versatile tools for programmable gene regulation and synthetic biology. In its catalytically inactive form (dead Cas9, dCas9), the protein retains its RNA-guided DNA-binding capacity but lacks nuclease activity, allowing it to function as a programmable transcriptional regulator (see Fig. 1-A, top). By coupling specific single guide RNAs (sgRNAs) with dCas9, transcriptional repression (CRISPR interference, CRISPRi, Fig. 1-B, top) or activation (CRISPRa, Fig. 1-B, bottom) can be achieved at virtually any genomic locus containing a protospacer adjacent motif (PAM) sequence [1]. This programmability and orthogonality make CRISPR an ideal foundation for constructing genetic circuits.

**Fig. 1.**
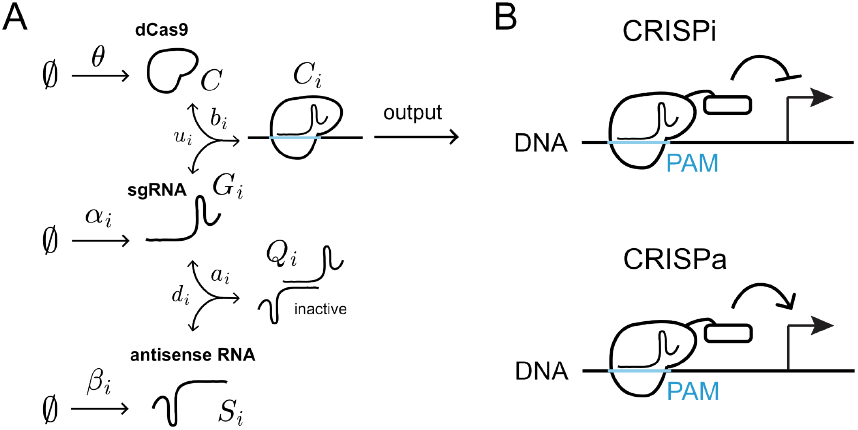
Design of a CRISPR-based perceptron. Panel A shows the schematic of the chemical reaction network underlying the circuit, including dCas9, guide RNAs, antisense guide RNAs, and their binding interactions leading to formation of regulatory dCas9–gRNA complexes that target PAM-containing DNA sites. Once bound to DNA, the dCas9–gRNA complex mediates either transcriptional repression (top, CRISPRi) or transcriptional activation (bottom, CRISPRa), depending on the effector domain fused to dCas9, as illustrated in Panel B.

Another key advantage of the CRISPR-dCas9 platform is that regulatory domains can be modularly fused to dCas9, enabling gRNA-specified activation or repression, which enables coupling between a decision-making circuit to a transcriptional response. This makes CRISPR an attractive substrate for implementing molecular classifiers [2], [3]. Here, we propose a novel architecture for biomolecular neural networks constructed from the dCas9 protein, guide RNAs, and antisense RNAs. In our design, antisense RNA is used to sequester sgRNAs through the formation of inactive antisense RNA-sgRNA complexes [4]. This molecular titration mechanism enables precise control over the pool of functional sgRNAs available for dCas9 binding, providing a tunable regulatory layer within the circuit. Using mathematical and computational analysis, we show that this system functions as a perceptron at steady state. However, extending a single perceptron into a multilayer architecture creates competition for a finite pool of dCas9, whose expression must remain low to avoid cytotoxicity [5]. To address this limitation, we develop a resource-aware design framework for CRISPR-based molecular classifiers and demonstrate its potential for executing both classification and regression tasks.

## II. Design of a CRISPR-based perceptron

### A. Modeling competition for the dCas9 protein

In the minimal implementation of the CRISPR-based perceptron, the inactive dCas9 protein (*C*) is constitutively produced and degraded at rates *θ* and δ, respectively. The guide RNA (*G*_*i*_), produced at rate constant *α*_*i*_, reversibly binds dCas9 at rate *b*_*i*_ to form the active CRISPR complex (*C*_*i*_), which dissociates at rate *u*_*i*_. The antisense RNA (*S*_*i*_), produced at rate constant *β*_*i*_, reversibly binds *G*_*i*_ at rate *a*_*i*_, sequestering it and preventing association with dCas9 by forming an RNA complex (*Q*_*i*_) that dissociates at rate *d*_*i*_. The index *i* denotes the number of sgRNA species in the system, and *i* = 1 corresponds to the minimal perceptron architecture schematized in Fig. 1-A.

Using this notation, we generalize the architecture to a multi-sgRNA CRISPR network, in which each sgRNA species can form its own dCas9–sgRNA complex and all species compete for the same finite pool of dCas9. Under this shared-resource regime, the CRISPR-based perceptron is described by the following chemical reactions:

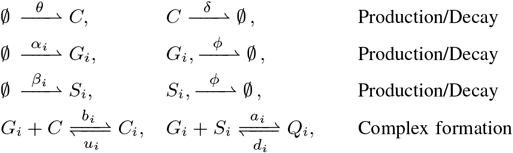

for *i* = 1, …, *m*. Using the law of mass action, we can model the dynamics of each chemical species’ concentration using the following Ordinary Differential Equations (ODEs):

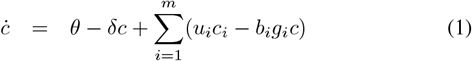

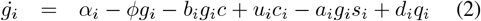

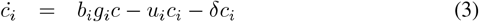

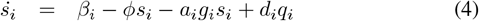

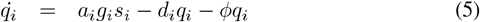

### B. Steady-state analysis

We state the mass conservation equation for the dCas9 protein by defining its total concentration as the combined amount of unbound (free) dCas9 and dCas9 complexed with sgRNA, resulting in the following expression, obtained

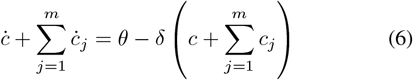

Solving for the steady-state value of the dCas9 protein variable *c*(*t*), denoted as 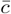, we equate (6) to zero, resulting in

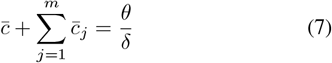

Similarly, solving for the steady state of the dCas9–sgRNA complex variable *c*_*i*_(*t*), denoted by 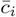, we equate (3) to zero, resulting in

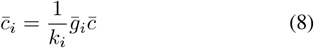

where *k*_*i*_ = (*u*_*i*_ + δ)*/b*_*i*_. Then, by substituting Eq. (8) into Eq. (7), we obtain

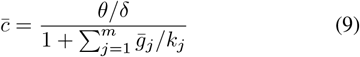

Replacing Eq. (9) into Eq. (8) yields

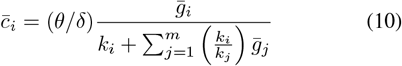

Moreover, solving for the steady state of the inactive RNA complex variable *q*_*i*_(*t*), denoted by 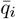, we equate (5) to zero, resulting in

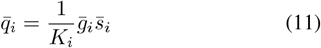

where *K*_*i*_ = (*d*_*i*_ + *ϕ*)*/a*_*i*_. Then, solving for the steady state of the antisense RNA variable *s*_*i*_(*t*), denoted by 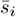, we equate (4) to zero:

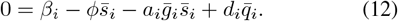

Replacing Eq. (11) in Eq. (12), and solving for 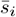 results in

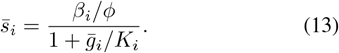

Furthermore, replacing 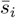 back into Eq. (11) gives

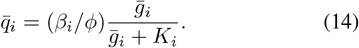

Using the steady-state relations 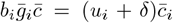 from Eq. (3) and 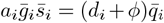 from Eq. (5), we can rewrite the steady-state expression for Eq. (2) as follows:

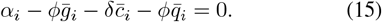

Substituting 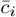 from Eq. (10) and 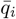 from Eq. (14) into Eq. (15) yields

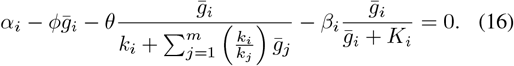

For convenience, let 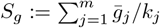. Using Eq. (10) for 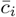 and factoring *k*_*i*_ from the denominator, we have

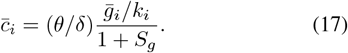

Similarly, let 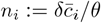. Then Eq. (17) gives

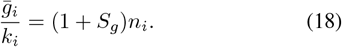

Summing over *i* = 1, …, *m* gives

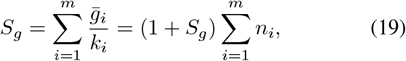

which can be solved for *S*_*g*_:

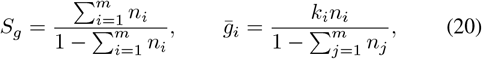

valid for 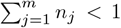. Finally, substituting Eq. (20) into Eq. (16) and dividing by *θ* gives the following nondimensional equation

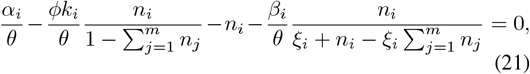

where *ξ*_*i*_ = *K*_*i*_*/k*_*i*_. Since our goal is to determine the closed-loop steady-state solution for 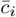 to demonstrate its behavior as a perceptron, in the following sections we analyze the solutions of Eq. (21).

### C. Approximated input-output response in the fast sequestration regime

Let *N*_*i*_ describe the free (unbound) dCas9 protein concentration, so that

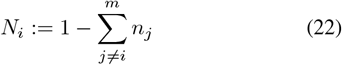

Replacing in Eq. (21), we obtain

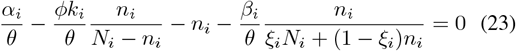

For *N*_*i*_ *>* 0, we introduce the following non-dimensionalized representations: 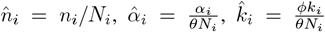, and 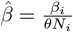. Then, we can write a non-dimensionalized version of Eq. (23) as follows:

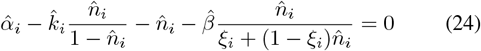

Solving for 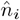 gives a polynomial equation of the form 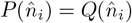, where

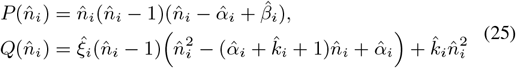

To understand whether the roots converge we numerically solved 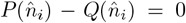 in the limit 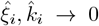 and examined the three parameter regimes 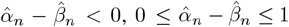, and 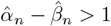. The three roots collapse to 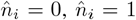, and 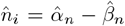 in that limit, which of these is physically feasible (i.e., that lies inside 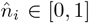, shown in gray in Fig. 2) depends on the (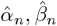) regime. In particular,

**Fig. 2.**
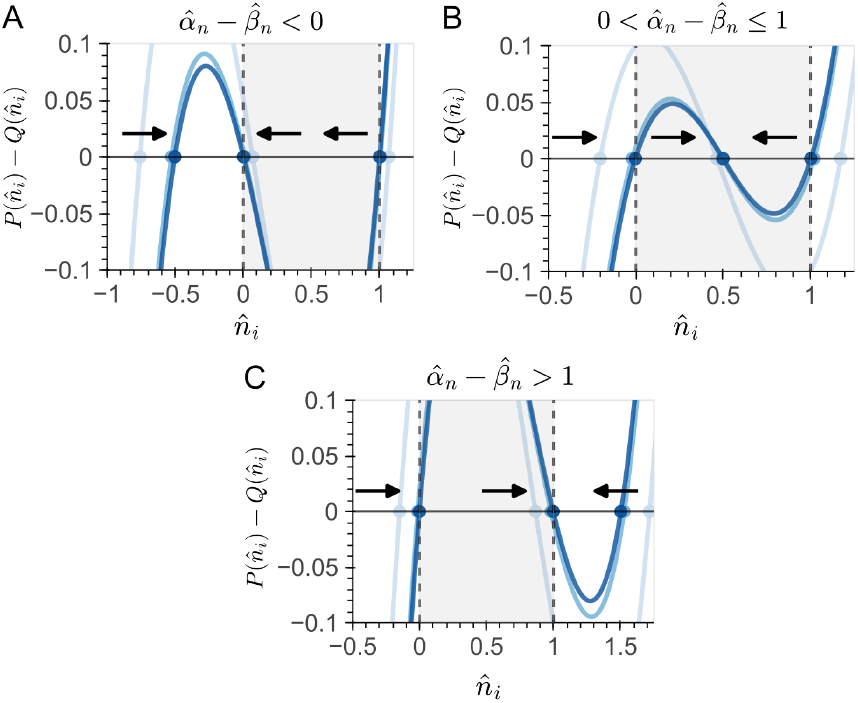
Root analysis in the fast sequestration regime. Panels A–C show 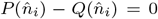 computed for three representative choices of 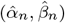 corresponding to the regimes 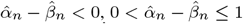, and 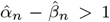. For illustration, we used 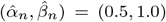in Panel A, (1.0, 0.5) in Panel B, and (2.0, 0.5) in Panel C. Curves show the resulting values of 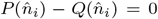, with filled circles marking the locations to which the three roots approach as 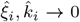 (from light to dark blue). Arrows indicate the direction of convergence along the 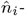 axis in each regime, and the gray highlighted area represents the physically admissible bounds for 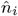.

- In Fig. 2-A, where 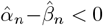, the interior root 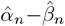 lies below the interval [0,1], while the 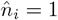 root lies just above it. Thus, the only admissible steady state is 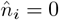. The entry 0^+^ in Table I indicates that this state is approached from the right as the small parameters vanish, consistent with the flow arrows in the figure.
- In Fig. 2-B, where 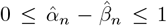, the nontrivial root 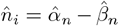lies inside [0,1] and is therefore the unique physically relevant solution. The roots at 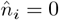and 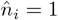 lie just outside the interval and appear as 0^−^ and 1^+^ in Table I. The trajectories in the figure converge toward the interior root.
- In Fig. 2-C, where 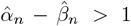, the interior root exceeds the upper limit of the interval, while 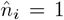 lies inside it. This admissible root is approached from the left, recorded as 1^−^ in Table I.

Thus in each regime there is a single physically feasible limit for 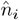 (respectively 0, 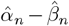, or 1), and the superscripts in Table 1 indicate the side from which the boundary root is approached in the 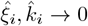 limit. To formalize the root analysis, we perform local expansions in the limit 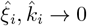 and relate them to the asymptotic behavior summarized in Table I.

**TABLE 1.**
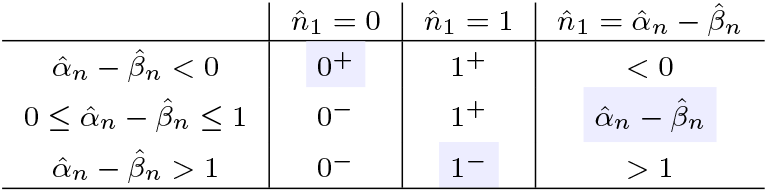
Summary of asymptotic root locations.

| | $\hat{n}_1 = 0$ | $\hat{n}_1 = 1$ | $\hat{n}_1 = \hat{\alpha}_n - \hat{\beta}_n$ |
| --- | --- | --- | --- |
| $\hat{\alpha}_n - \hat{\beta}_n < 0$ | $0^+$ | $1^+$ | $< 0$ |
| $0 \leq \hat{\alpha}_n - \hat{\beta}_n \leq 1$ | $0^-$ | $1^+$ | $\hat{\alpha}_n - \hat{\beta}_n$ |
| $\hat{\alpha}_n - \hat{\beta}_n > 1$ | $0^-$ | $1^-$ | $> 1$ |

First, we expand near equilibrium 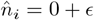, where *ϵ* is a small parameter ( |*ϵ*| ≪1). Keeping only the lowest-order terms in the left- and right-hand sides of Eq. (25) gives

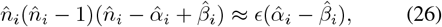

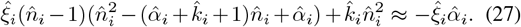

Balancing left and right sides, we find

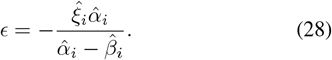

Hence, for small 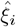, the root at 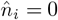 approaches from the left if 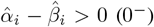 and from the right if 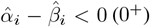, consistent with the first column of Table I.

Next, we expand near equilibrium 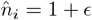, again with |*ϵ*| ≪ 1. Expanding to first order gives

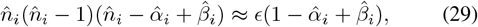

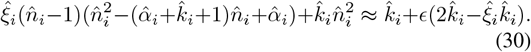

Balancing terms, we obtain

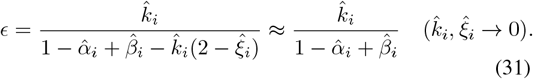

Thus, for small positive 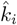, the root at 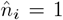 approaches from the right if 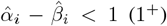 and from the left if 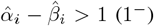, matching the second column of Table I. Ultimately, the third root, 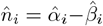, remains at its nominal location and lies within [0,1] whenever 0 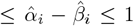, reproducing the third column of Table I. Consequently, we can approximate the only admissible solution of Eq. (24) as follows

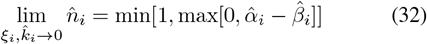

## III. How does shared dCas9 affect neuromorphic computation?

### A. Input-output mapping under competition for dCas9

Eq. (32) describes the steady-state activation function generated by the CRISPR-based perceptron and exhibits two distinct regimes. The first is a *threshold regime*, determined by the sign of the 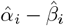 term. This condition specifies the sign of the weight associated with each input: an input that increases sgRNA production is assigned a positive weight, whereas an input that increases antisense RNA production is assigned a negative weight. Once the threshold is crossed, the activation increases linearly with the difference 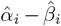. The second regime is *saturation*, imposed by the finite availability of the shared dCas9 pool through the min[·] operator.

Since 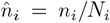, and 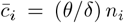, we can rewrite Eq. (32) as

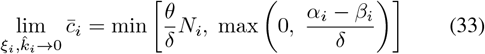

Eq. (33) shows that *N*_*i*_ directly rescales the saturation regime: as competing sgRNAs sequester dCas9 (reducing *N*_*i*_), the maximum achievable output decreases proportionally, rescaling the activation function. However, the threshold condition *α*_*i*_ = *β*_*i*_ is maintained.

To characterize how this approximation relates to the full reaction network, we numerically integrated Eqs. (1)–(5) for *i* = 1 while decreasing the parameter *ξ*_1_ = *k*_1_*/K*_1_, which quantifies the ratio between the formation rate of the sgRNA–dCas9 complex and the formation rate of the sgRNA–antisense RNA complex. Here, we explicitly define the inputs *X*_1_ and *X*_2_ producing the sgRNA and antisense RNA molecules at rate *w*_1_ and *w*_2_, respectively. As *ξ*_1_ decreases, shown in Fig. 3-B, the steady-state input–output curve displays a sharper threshold, the linear region, and a saturation plateau, as approximated by Eq. (32). Then, under non-saturating conditions and for a sufficiently small *ξ*_1_, the activation function for the CRISPR system resembles a saturated Rectified Linear Unit (ReLU) activation function [6].

**Fig. 3.**
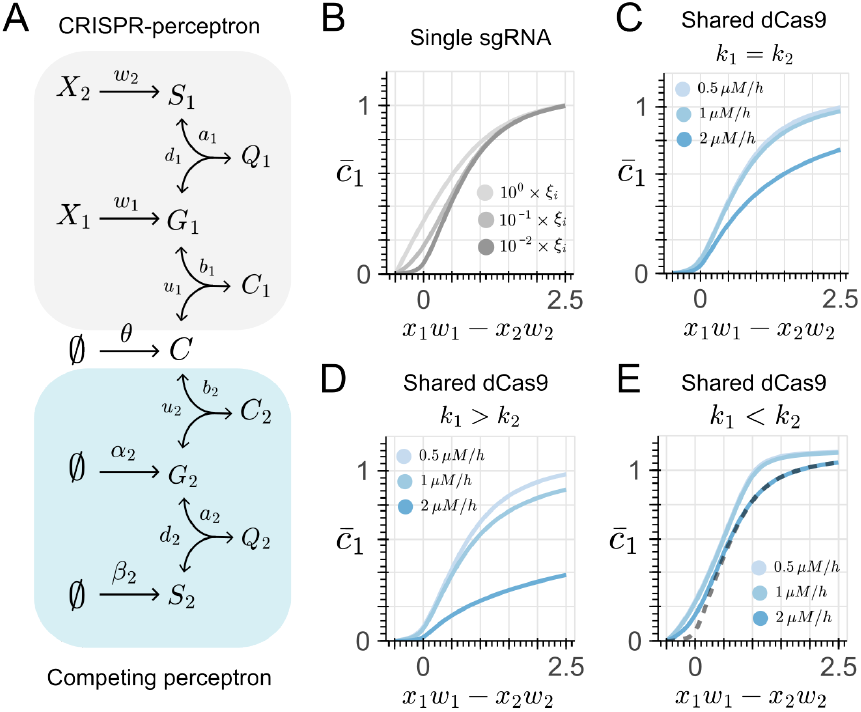
Effect of shared dCas9 protein in the activation function of the CRISPR-based perceptron. Panel A shows a schematic of the chemical reactions for two perceptrons sharing dCas9 resources, with the analyzed perceptron in gray and the competing one in blue. Panel B illustrates how the ratio between the sgRNA-dCas9 complex formation rate and the sequestration rate influences the input–output relationship: when *ξ* → 0 (*K*_*i*_ ≫ *k*_*i*_; strong RNA sequestration relative to dCas9 binding), Eq. (32) simplifies to a linear relationship, where the output is proportional to the input below saturation and plateaus at the capacity of the dCas9 pool at saturation. Panels C–E show the effect of varying dCas9 binding affinities on the output: Panel C corresponds to equal affinities (*k*_1_ = *k*_2_), Panel D to a higher affinity for the competing perceptron (*k*_1_ *> k*_2_), and Panel E to a higher affinity for the perceptron of interest (*k*_1_ *< k*_2_). All simulations used the following nominal kinetic parameters: *θ* = 1 *µM/h*, δ = *ϕ* = 1 */h, b*_1_ = *b*_2_ = 10 */µM/h, u*_1_ = *u*_2_ = 1 */h, d*_1_ = *d*_2_ = 1 */h, a*_1_ = *a*_2_ = 500 */µM/h, w*_1_ = *w*_2_ = 1 */h* and *β*_2_ = 1 *µM/h*. The input *x*_1_ was varied from 0 *µM* to 3 *µM*, while *x*_2_ was fixed at 0.5 *µM*. To simulate higher binding affinity for the dCas9 protein, the *b*_*i*_ parameter was increased 10-fold. For the competing perceptron, *α*_2_ was varied between 0.5, 1.0, and 2.0 *µM/h* to generate the dCas9 demand values (*C*^∗^) shown in the legends.

We next examined how competition for dCas9 modifies this mapping by introducing a second CRISPR perceptron producing an additional sgRNA/antisense pair (*G*_2_, *S*_2_) that shares the same dCas9 pool (Fig. 3-A), simulated by numerically integrating Eqs. (1)–(5) for *i* = 2 (see Appendix). Increasing the expression of the competing sgRNA reduces the magnitude of the activation of the first perceptron, while the threshold remains unchanged (Fig. 3-C), as described by Eq. (33). When the competing sgRNA has higher affinity for dCas9, the reduction in magnitude is greater (Fig. 3-D), whereas higher affinity for the sgRNA of interest makes the activation more robust (Fig. 3-E). Intuitively, both increased production of the competing sgRNA and increased affinity to the shared dCas9 pool lead to greater sequestration of the limiting dCas9 resource, thereby reducing the amount of dCas9–sgRNA complex available for the perceptron of interest while leaving the threshold condition (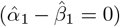) unchanged.

### B. Regression under competition for dCas9

From a systems perspective, endogenous signaling can be understood as a process that maps specific inputs, such as ligands, to downstream transcriptomic responses [7], [8]. By examining the input-output relationship, we can effectively capture the function that describes this mapping. In a synthetic biology context, this regulatory function can be reconstructed using genetic circuits, allowing us to build synthetic analogues of natural regulatory programs [9]. Here, we explore the application of a CRISPR-based network to approximate the input-output map of an endogenous process, which can be framed as a regression task. As a proof of principle, we focus on approximating a band-pass function, which is characteristic of certain developmental programs [10], [11].

In Fig. 4-A (top), we show the architecture for a single sgRNA regressor. In this design, *x* represents the input that drives sgRNA production, while *T*_*i*_ (*i* = 1, …, *n*) is a fixed input producing antisense RNA that sets the threshold described in Eq. (32). Building on this single-node design, we propose an multi-layer architecture (MLP; Fig. 4-A, bottom) to reconstruct the band-pass function (see Appendix). To understand how the MLP approximates the target function, we visualize the output of each individual perceptron. Specifically, Fig. 4-A1 shows the input-output map for Node 1, with threshold set at *x* = 1, *µ*M, while Fig. 4-A2 shows the map for Node 2, with threshold at *x* = 2, *µ*M. The combined output of the MLP, shown in Fig. 4-A3, resembles the band-pass function. Furthermore, the tunability of this architecture is illustrated in Fig. 4-A4: adjusting the thresholds for Node 1 and Node 2 produces a narrower band-pass.

**Fig. 4.**
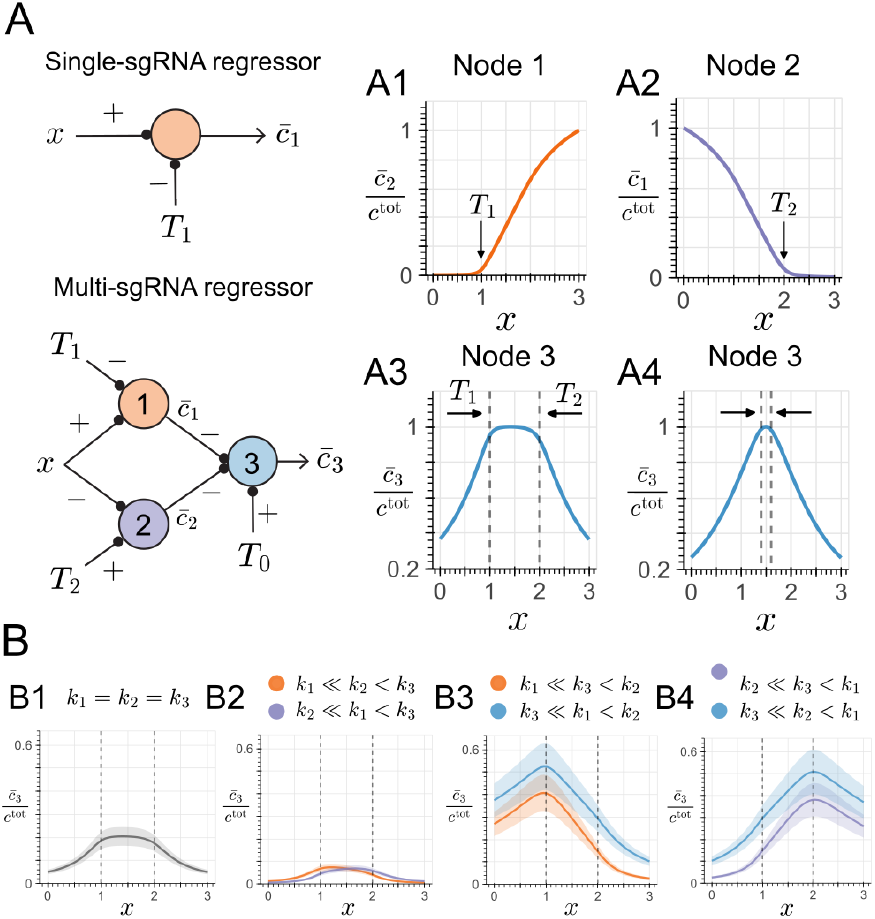
Design of a CRISPR-based regressor under competition for dCas9 protein. Panel A (left) illustrates the design of CRISPR-based regressors for a single sgRNA (top) and multiple sgRNAs (bottom), the latter equivalent to an MLP with two nodes in the hidden layer and one node in the output layer. Following Eq. 32, *x* represents the input, while *T*_*i*_ (*i* = 1, 2, …) represents a fixed input concentration that sets the threshold. Panels A1–A3 show the output of each node of the MLP, color-coded by node: orange for Node 1 (*T*_1_ = 1, *µ*M), purple for Node 2 (*T*_2_ = 2, *µ*M), and blue for Node 3, which approximates the band-pass function. Panel A4 shows the same output for a narrower threshold range (*T*_1_ = 1.4, *µ*M, *T*_2_ = 1.6, *µ*M). Panel B illustrates the effect of varying dCas9 binding affinities on the output: Panel B1 corresponds to equal affinities, Panel B2 to the sgRNA in Node 3 having the smallest affinity, Panel B3 to Node 2 having the smallest affinity, and Panel B4 to Node 1 having the smallest affinity. All simulations used the same kinetic parameters and binding affinities listed in Fig. 5, and the uncertainty regions in Panel B were defined by varying the binding affinities by *±*20% of their nominal values.

Since each node in the MLP competes for a finite pool of dCas9 protien, to explore the impact of this competition, in Fig. 4-B we systematically screen all combinations of heterogeneous binding affinities between the sgRNAs in each node and dCas9. As observed previously, the magnitude of the MLP output is sensitive to these affinity combinations, as described by Eq. (33). For instance, compared to the equal-affinity baseline in Fig. 4-B1, when the output node has the smallest sgRNA affinity, the output magnitude is roughly halved (Fig. 4-B2), whereas when the output node has the highest affinity, the output magnitude increases. In all cases, the thresholds remain unchanged, but the shape of the band-pass function varies with binding affinity. Note that every regressor in Fig. 4-B include uncertainty regions to illustrate context-dependent variation in sgRNA-dCas9 binding affinity *in vivo*.

### C. Classification under competition for dCas9

Given that sharing the dCas9 pool does not alter the thresholding behavior of the CRISPR perceptron, we expect the decision boundary of a linear classifier to remain insensitive to this effect. Fig. 5-A shows an abstract representation of the reaction network, where the signs are given by the max(·) operator of Eq. (32). The heatmap in the *x*_1_ − *x*_2_ plane in Fig. 5-A1 corresponds to a perceptron without competition and exhibits a linear decision boundary. Introducing a competing sgRNA controlled by the rate *α*_2_ decreases the magnitude of the output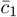, but the location and orientation of the decision boundary in the *x*_1_ − *x*_2_ plane remain unchanged.

**Fig. 5.**
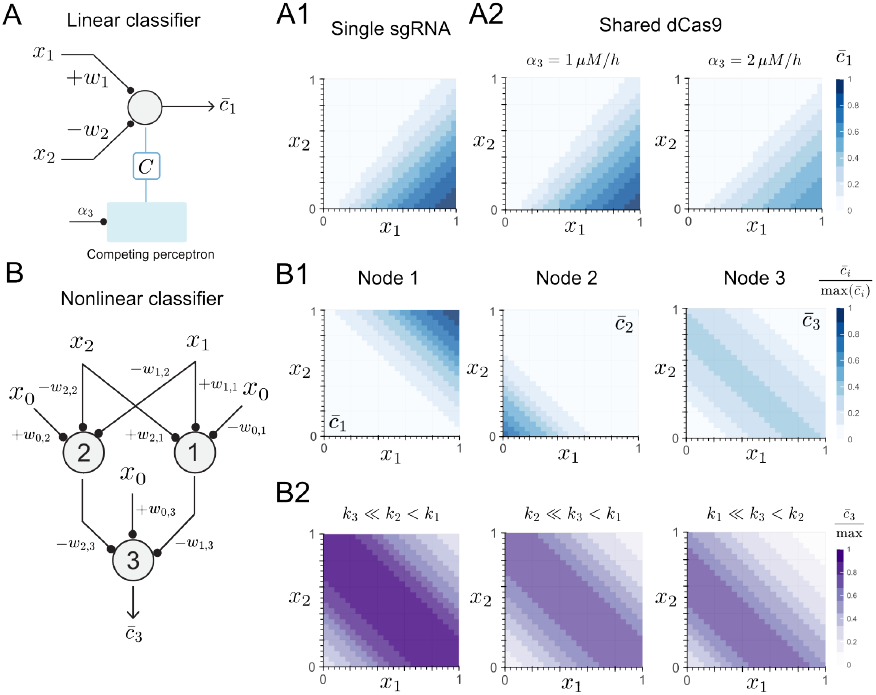
Linear and nonlinear CRISPR-based classifiers under competition for dCas9 protein. Panel A illustrates the perceptron abstraction for the chemical reaction network introduced in Fig. 3, and shows the linear decision boundaries for a single sgRNA (Panel A1, without competition), and for the shared dCas9 protein setup (Panel A2). Similarly, Panel B illustrates the multi-layer perceptron (MLP) consisting of 2 nodes in the hidden layer and 1 node in the output layer. Panel B1 shows the linear decision boundaries generated by Node 1 and Node 2, as well as the nonlinear decision boundary generated by Node 3. Panel B2 shows the nonlinear decision boundary generated by Node 3 under diferent binding affinity combinations between the sgRNA of each node and the dCas9 protein. The heatmaps in the *x*_1_ − *x*_2_ plane in Panels A1 and A2; B1 and B2 were normalized by the maximum value for the single sgRNA simulation, by the maximum value across the nodes, and by the maximum value for the highest affinity for the sgRNA in Node 2, respectively. The same kinetic parameters listed in Fig. 3 were used in all simulations, except for the binding affinities in Panel B2, which were computed from the set *b*_*i*_ = (10, 50, 100) */µM/h*, for *i* = {1, 2, 3}.

To explore how this effect propagates in larger networks, we next consider a an MLP with two hidden nodes and one output node that all draw from the same finite dCas9 pool, with a similar design to that shown in Fig. 4-A. The nonlinear decision boundary produced by the output node originates from the composition of the two linear boundaries defined by Nodes 1 and 2, as illustrated in Fig. 5-B1 (see Appendix). When all sgRNAs have equal affinity for dCas9, only the output node exhibits a reduced magnitude. This difference follows from the network architecture: the hiddennodes sequester dCas9 before the forward pass, leaving a smaller fraction of the protein available to the final node. Nonetheless, this effect is only at scale of the magnitude: the characteristic band-shaped classification region remains clearly defined.

Moreover, for heterogeneous affinities, as shown in Fig. 5-B2, increasing the affinity of the sgRNA of the output node allows it to sequester more dCas9, which increases its output magnitude relative to the equal-affinity case in Fig. 5-B1. In contrast, changing the affinities of the sgRNAs of the hidden nodes shifts their linear boundaries in the *x*_1_− *x*_2_ plane. For example, higher affinity of the sgRNA in Node 2 moves its linear decision boundary toward the top-right, whereas an higher affinity sgRNA in Node 1 moves its linear decision boundary toward the bottom-left. These shifts reposition the nonlinear band-shaped classification region (see shifts, from left to right, in Fig. 5-B2) and reduce the magnitude of the output node. Despite these variations, the band-like classification region remains clearly defined across all affinity combinations.

## IV. Discussion

Here, we demonstrated that CRISPR-dCas9 systems with antisense RNA sequestration can function as molecular perceptrons capable of both classification and regression tasks. Our analysis reveals that these circuits generate thresholding activation function that resemble a saturated ReLU in the fast sequestration regimen. Moreover, through numerical simulations, we found that competition for shared dCas9 resources affect output magnitude but preserve the underlying logic. For classification, decision boundaries remain invariant under resource-constrained conditions, while for regression, the thresholds set by individual nodes remain robust to resource competition, although the output magnitude shows sensitivity to heterogeneous binding affinities.

This work expands the repertoire of molecular substrates available for neuromorphic computing in living cells. Alongside existing approaches based on molecular sequestration [12], [13], phosphorylation/dephosphorylation cycles [14], and enzymatic reactions [15], CRISPR-based circuits offer unique advantages in programmability and modularity. Beyond solving classification tasks, the potential to approximate biologically relevant regulatory functions (provided that the input-phenotype mapping is known) suggests immediate applications reconstructing and controlling endogenous signaling programs [9]. Ultimately, by establishing a resource-aware design framework that explicitly accounts for the finite availability of dCas9, we provide a foundation for engineering more complex multilayer architectures that can execute highly complex computation tasks within the constraints of cellular biochemistry.

## V. Appendix A. CRISPR perceptron under shared dCas9

The full model of the CRISPR perceptron, along with the competing circuit shown in Fig. 3-A, is given by the following chemical reactions:

1. Production and decay of dCas9 pool

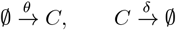
2. *2) CRISPR-based perceptron (node of interest)*

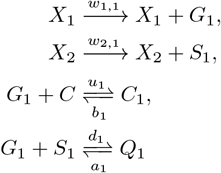
3. *3) Competing circuit*

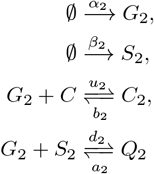

## APPENDIX B Ordinary Differential Equations (ODEs)

The chemical reactions stated in the previous subsections can be modeled by the following ODEs:

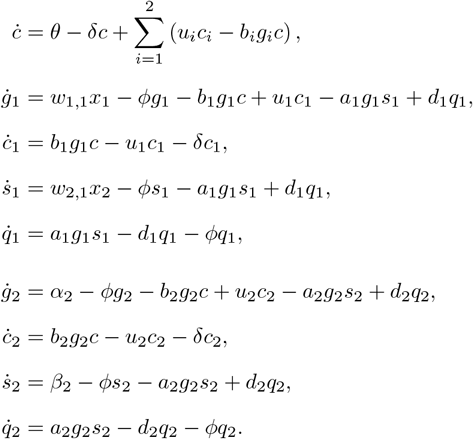

## APPENDIX C Multi-sgRNA regressor

The full model of the MLP shown in Fig. 4-A (bottom) is given by the following chemical reactions:

1. *Production and decay of dCas9 pool*

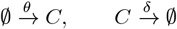
2. *Node 1 (hidden layer)*

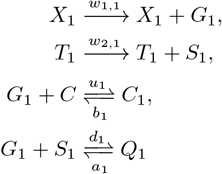
3. *Node 2 (hidden layer)*

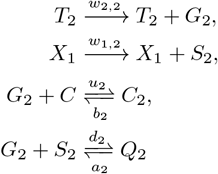
4. *Node 3 (output layer)*

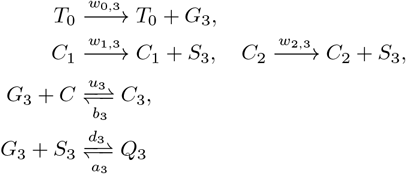
5. *Ordinary Differential Equations (ODEs)* The chemical reactions stated in the previous subsections can be modeled by the following ODEs:

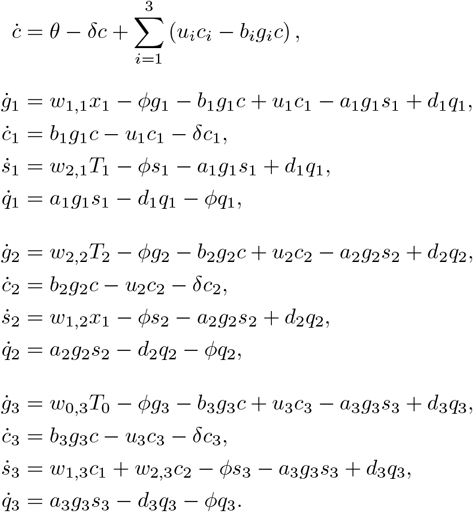

## APPENDIX D Multi-sgRNA classifier

The full model of the MLP shown in Fig. 5-B is given by the following chemical reactions:

1. *Production and decay of dCas9 pool*

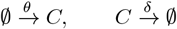
2. *Node 1 (hidden layer)*

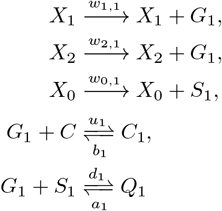
3. *Node 2 (hidden layer)*

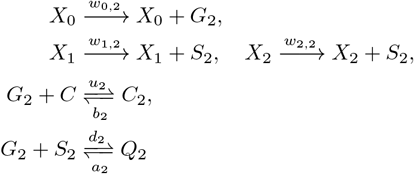
4. *Node 3 (output layer)*

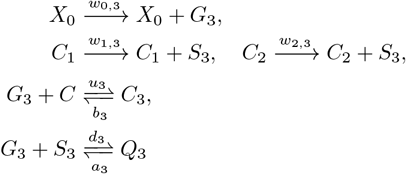
5. *Ordinary Differential Equations (ODEs)* The chemical reactions stated in the previous subsections can be modeled by the following ODEs:

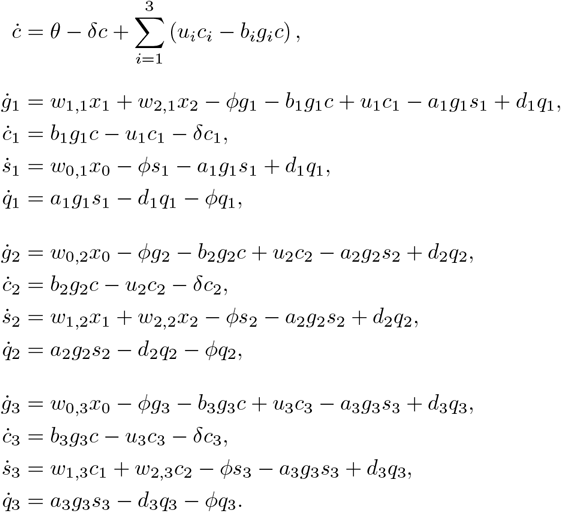

## Notes

### Competing Interest Statement

The authors have declared no competing interest.

